# Ontology-based Protein-Protein Interaction Explanation Using Large Language Models

**DOI:** 10.1101/2025.04.07.647599

**Authors:** Nur Bengisu Çam, Hasin Rehana, Jie Zheng, Benu Bansal, Yongqun He, Junguk Hur, Arzucan Özgür

**Author notes:** Co-first authors (Nur Bengisu Çam, Hasin Rehana).

## Abstract

Protein-protein interactions (PPIs) play a crucial role in various biological processes, and understanding these interactions is essential for advancing biomedical research. Automated extraction and analysis of PPI information from the rapidly growing scientific literature remains an important challenge. We present a novel ontology-based approach to analyze protein-protein interactions using Large Language Models (LLMs). We applied different learning strategies, namely in-context learning and parameter-efficient instruction fine-tuning for the Llama-2 chat models, to identify keywords in the text that indicate an interaction between a pair of proteins. Our results show that parameter-efficient fine-tuning leads to a performance gain even when the domain is new. The smaller fine-tuned models outperformed the zero-shot performance of much larger models. The keywords identified by the Llama-2 models were mapped to the ontology terms in the Interaction Network Ontology (INO). Our study suggests that a pipeline of an LLM and an ontology is an effective strategy for explaining relations between biomedical entities. This work demonstrates the potential of leveraging ontologies and advanced language models to advance automated PPI analysis from the scientific literature.

## I. Introduction

In biomedical text mining, the relations between biomedical entities are expressed by words, which can be called as relation keywords. These relation keywords can be used to infer the type of the relation between pairs of entities. In our previous study, we collected 826 interaction keywords (such as binds, bound, interact, activates, and phosphorylates), and the usage of these keywords supports our literature-based discovery of IFN-gamma and vaccine-mediated gene interaction networks [1]. Note that some of these keywords are variations of the same words (e.g., binds and bound). These variations of the same words were also compiled and included in our list of collected keywords.

Furthermore, we realized that these keywords share semantic relations. For example, phosphorylation is a special type of chemical interaction, and ‘phosphorylates’ is a variation of representing the phosphorylation interaction. To better support literature mining of these keywords, we developed the Interaction Network Ontology (INO) and used INO to represent various types of interactions [2]–[4].

In this study, we present an approach to explain the relations between proteins described in biomedical text by identifying the ontological keywords for interactions in the corresponding sentences. Here we harnessed the power of large language models (LLMs), in particular the Llama-2 chat models. Our methods include contextual learning with zero-shot and fewshot settings, as well as parameter-efficient instruction finetuning. We compared the models using these learning techniques. Our goal was to show that the models with smaller parameters, when fine-tuned even with parameter-efficient techniques, can outperform the larger models. Here, the finetuned Llama-2-13b-chat model was superior to the Llama-2-70b-chat model with few-shot learning.

## II. Related Work

Following the development of the transformer architecture, the era of LLMs has begun. The LLMs are known for their large corpus pre-training, enormous parameter size, and massive data processing capabilities for identifying connections in text elements. Therefore, the pre-training and subsequent finetuning of language models became popular for various NLP domains, such as biomedical text mining.

Bidirectional Encoder Representations from Transformers (BERT) is an encoder-based LLM trained on the general corpora. Various BERT-based models have emerged and become popular for biomedical text mining. BioBERT [5], Bidirectional Encoder Representations from Transformers for Biomedical Text Mining, is an LLM specifically pre-trained on biomedical text from PubMed articles. ClinicalBERT [6] further pre-trained the BERT model for comparing the effect of domain-specific contextual embeddings. SciBERT [7], on the other hand, is an LLM pre-trained on scientific texts that also contain biomedical data. It has been shown that pre-training with biomedical text shows superior performance on domain-specific tasks [5], [7] compared to pre-training with general domain information.

Generative pre-trained transformer (GPT) is a decoder-based LLM that generates human-like text through the acquisition of linguistic patterns and structures [8]. On the other hand, ChatGPT is a GPT-based chat model developed by OpenAI [9]. It is a natural and engaging conversation tool that can produce contextually relevant responses based on text data. In our previous study [10], we evaluated the zero-shot performance of ChatGPT and compared the results with BioBERT, SciBERT, and PubMedBERT on protein-protein interaction tasks using the LLL, HPRD50, and IEPA datasets. The results showed that while ChatGPT performed competitively with carefully designed prompts, the BERT-based models were superior.

Recent studies have investigated the performance of LLMs in zero-shot and few-shot learning for various tasks such as analyzing, producing, and understanding text [11], [12]. Zeroshot and few-shot performances were reported to outperform fine-tuned models on question-answer datasets [11].

On the other hand, the advantages of parameter-efficient fine-tuning (PEFT) over in-context learning were investigated on a small dataset [13]. It was shown that their PEFT method not only outperforms the results of in-context learning but is also cheaper in terms of computational costs on the Real-world Annotated Few-shot Tasks (RAFT) benchmark [14].

Llama [15] is a decoder-based LLM developed by Meta. It was trained on a very large collection of corpora from general domains. One of the main advantages of Llama over GPT models is that Llama is completely open source. This allows researchers to make further improvements. It was reported that Llama performs better than GPT-3 on almost all benchmarks [15]. In the same year, the researchers developed a newer version of Llama, which is Llama-2.

There are domain-specific Llama models for biomedical text mining. One of them is PMC-Llama [16], which was developed specifically for medical applications, as the LLMs of the general domain lack medical information. The model was trained with a large unsupervised dataset consisting of 4.8 million articles and 30,000 books from the biomedical domain. The PMC-Llama model with 13 billion parameters outperformed the Llama-2 model with 70 billion parameters by 13.43% and the ChatGPT model by 9.46% on Q&A benchmarks.

Some other examples of biomedical Llama are MEDITRON-7B and MEDITRON-70B [17]. These are Llama-2-based models with 7 billion and 70 billion parameters, respectively, and were further pre-trained on carefully constructed medical corpora collected from PubMed Central and PubMed articles. It was reported that the in-context learning performance of MEDITRON-7B outperformed PMC-Llama by 10% on various biomedical benchmarks. Compared to the fine-tuned baseline models, MEDITRON-7B showed a performance gain of 5% and MEDITRON-70B showed a performance gain of 2%.

To our knowledge, we present here the first study demonstrating the effectiveness of an ontology- and LLM-based pipeline to explain relations between biomedical entities, in particular protein-protein interactions.

## III. Dataset

In our study, we used the Learning Language in Logic (LLL) INO dataset [4]. The LLL [18] focuses on extracting protein and gene interactions from biological abstracts related to Bacillus subtilis. On the other hand, LLL INO [4] contains the interaction sentences in the LLL dataset where gene/protein interactions are manually annotated with INO terms [3]. INO plays an important role in the logical representation of biological interactions, pathways, and networks. As of now, the INO ontology comprises a total of 575 terms, of which 202 terms fall into the interaction branch. The importance of the INO keyword representation is underlined by its ability to provide a structured set of ontological terms and associated keywords to support literature mining of genegene interactions in the biomedical literature. In many cases of biomedical relationships, two or more interaction keywords are used in combination to represent the complex interactions [4]. Therefore, the interactions in the LLL dataset were annotated with one or more INO terms. Figure 1 demonstrates the INO representation of ‘positive regulation’ and its relations with other interaction types^1^. Also, literature mining keywords are provided.

**Fig. 1.**
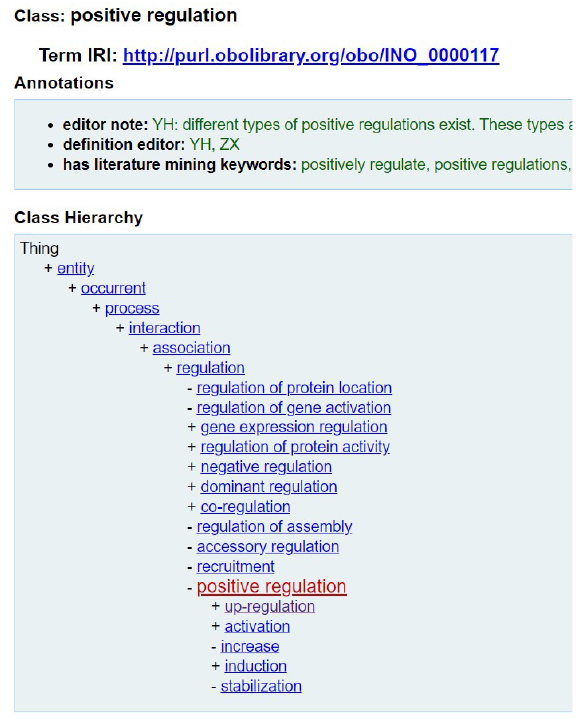
The INO representation of ‘positive regulation’ and its relations with other interaction types.

### A. Preprocessing

Before providing the examples into LLMs, we applied preprocessing. We followed a similar strategy to Li et al. (2021) [19] for processing the entities. Briefly, we enclosed the interacting protein names in the sentences with the tags [PROTEIN1]-[/PROTEIN1] and [PROTEIN2]-[/PROTEIN2] for a relation pair. We refer to this preprocessing step as protein tagging. It is helpful to indicate which proteins are interacting with each other, while keeping the protein or gene information as well.

We split the 164 total interaction sentences in the LLL INO dataset into three. After the split the training set has 125, the validation set has 14, and the test set has 25 interaction sentences.

## IV. Methods

We focused on explaining the relations by identifying relation keywords. Our goal was to find the relation keywords in a PPI sentence. We have used the LLL INO dataset [4]. This is the only available dataset that we can experiment with. Here, we chose Llama-2, a causal LLM, for several reasons. First, because Llama-2 has a decoder-transformer architecture like GPT and Llama language models are excellent for generation tasks due to their architecture and training properties [8].

For this task, we used in-context learning and parameter-efficient instruction tuning approaches on the LLL INO dataset. Of the various LLMs, Llama-2 variants were selected for this task because it has the same architecture as GPT-3, and the model is available as an open-source model.

### A. Updating INO

In our processing of the LLL dataset, which is tagged with INO terms, we identified some keywords that do not occur in INO. We handled those terms by comparing the lemma’s of the predicted and actual keywords. Eventually, we updated INO by adding these missing interaction keywords, including “action”, “inducible”, “production”, “assembly”, “transcriptional”, “act”, and “negatively controls”.

### B. In-Context Learning

#### 1) Zero-Shot Setting

In this setting, we experimented with zero-shot prompting for inferences with the Llama-2 chat models.

In our prompt, we have an introduction statement, an instruction statement, an input statement, and a response statement. The introduction statement consists of the introduction key and the introduction sentence. Next, we have the instruction statement. The previously mentioned studies have also experimented with similar instructions with chat models [20]. Previous experiments have shown that the instruction needs to be as clear and as informative as possible [10]. The instruction statement consists of the instruction key and the instruction sentence. The instruction sentence simply tells the model what to do for our specific task. The input statement consists of the input key and input sentence, where the input sentence is the test sentence we want to infer. Finally, the response statement consists of only a response key. This indicates that the model is required to respond after the response key. The prompt template for the zero-shot setting is shown in Figure 2. The prompt is created by combining the introduction, instruction, and input for each sentence in the test split of the LLL INO dataset.

**Fig. 2.**
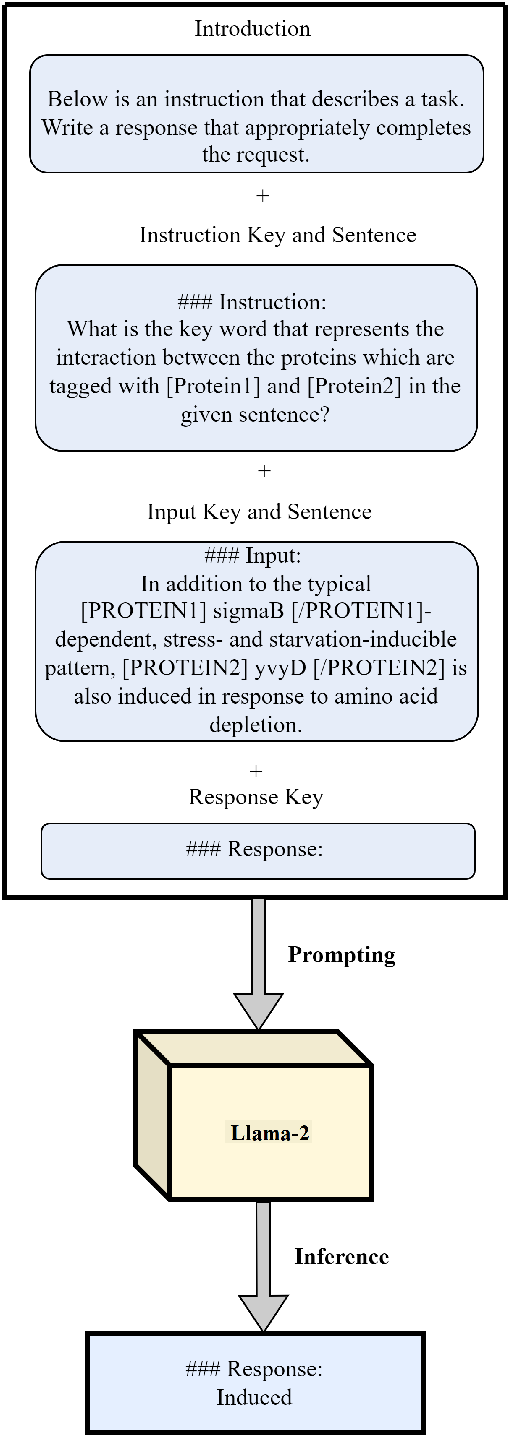
Zero-shot prompt example with a prompt that contains the introduction, instruction and input statements as well as a response key.

#### 2) Few-Shot Setting

The prompt template for the few-shot setting is shown in Figure 3. Here, we provide a series of examples in the prompt. This helps the model to learn from a small number of examples without the need for training or fine-tuning. With few-shot prompting, we take advantage of the power of in-context learning [11].

**Fig. 3.**
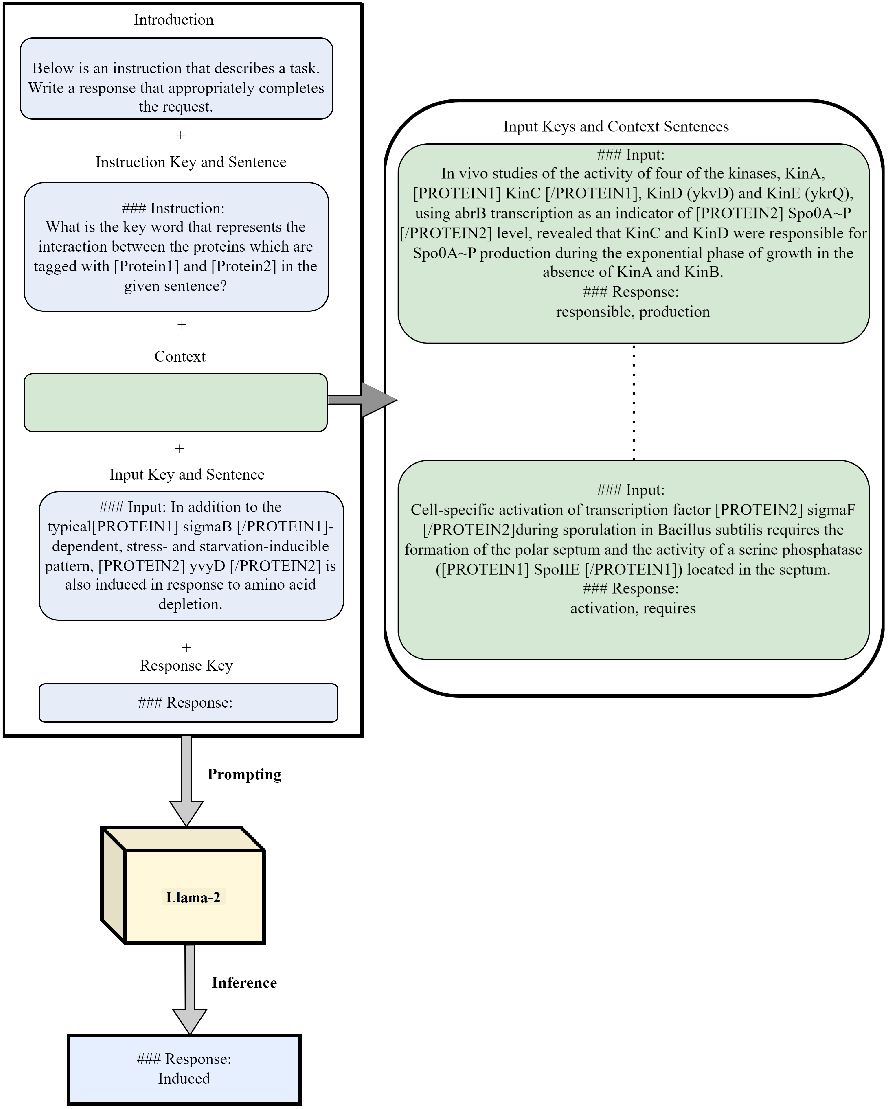
Few-shot prompt example with a prompt that contains the introduction, instruction, context, and input statements as well as a response key.

Here, we make the inference similar to the zero-shot prompting. The main difference is that we have a context with more than one input and response statement. In this context, several examples are listed one after the other with their consecutive INO terms. This is actually a list of input and response statements, where it stands as a series of examples for the given instruction statement. The model understands the task based on the instruction and the specified input-response pairs in the context. After the context, we have an additional input statement that has the test sentence. Finally, we have the response key in the response statement.

While selecting the number of examples in the context statement, we leveraged the long context of the Llama-2 models. Llama-2 [15] has a large context length of 4,096 tokens. This makes it possible to process more input tokens at once. In most cases, the more examples, the better the predictions [11]. Therefore, we have chosen the number of examples so that the context length is almost fully utilized. We calculated the average token length of the LLL INO sentences as around 80. We decided that the context should contain 40 sentence-response pairs and one test sentence. These prompt examples were selected from the training split of the LLL INO dataset.

In experiments with few-shot prompting, selecting the examples should be handled carefully. Depending on the representative examples provided in the context, the model may react differently. Therefore, we conducted three different fewshot experiments in which we ended up using almost all of the training examples. The overall performance of the few-shot prompting can be considered as the average score of these three experiments. Figure 3 illustrates the few-shot prompting example. The prompt is created by combining the introduction, instruction, context and input statements as well as a response key. This prompt style was constructed for each sentence in the test split of the LLL INO dataset.

### C. Parameter-Efficient Instruction Fine-tuning

Instruction fine-tuning is simply a fine-tuning technique where the inputs are instructions. Instructions are not only used for inference but can also be helpful in the training phase. Some studies have shown that model size may not be as important as we believe. Once the model is trained to follow the instructions, the results can outperform the performance of much larger LLMs [21]. Other studies revealed that if the model is instruction-tuned, the zero-shot performance increases on unseen tasks [22].

While fine-tuning, we used the 4-bit quantization technique, where the model is loaded with 4-bit precision. This configuration is available in the BitsAndBytes library [23]. In addition to quantization, we also used the LoRa configuration, which allows us to train only a small weight matrix [24]. This makes the training fast and memory efficient. We also used model parallelism to fit the models into the memory.

The prompt is similar to the zero-shot prompting structure described earlier. The main difference is in the response statement. Since the model needs to be trained using the training examples and consecutive INO terms, we provided the INO terms after the response key in the response statement. This prompt style was applied to every training example in the batch for several epochs. The next word prediction was applied, and the loss was back-propagated to update the model weights were updated. Figure 4 illustrates our instruction finetuning process.

**Fig. 4.**
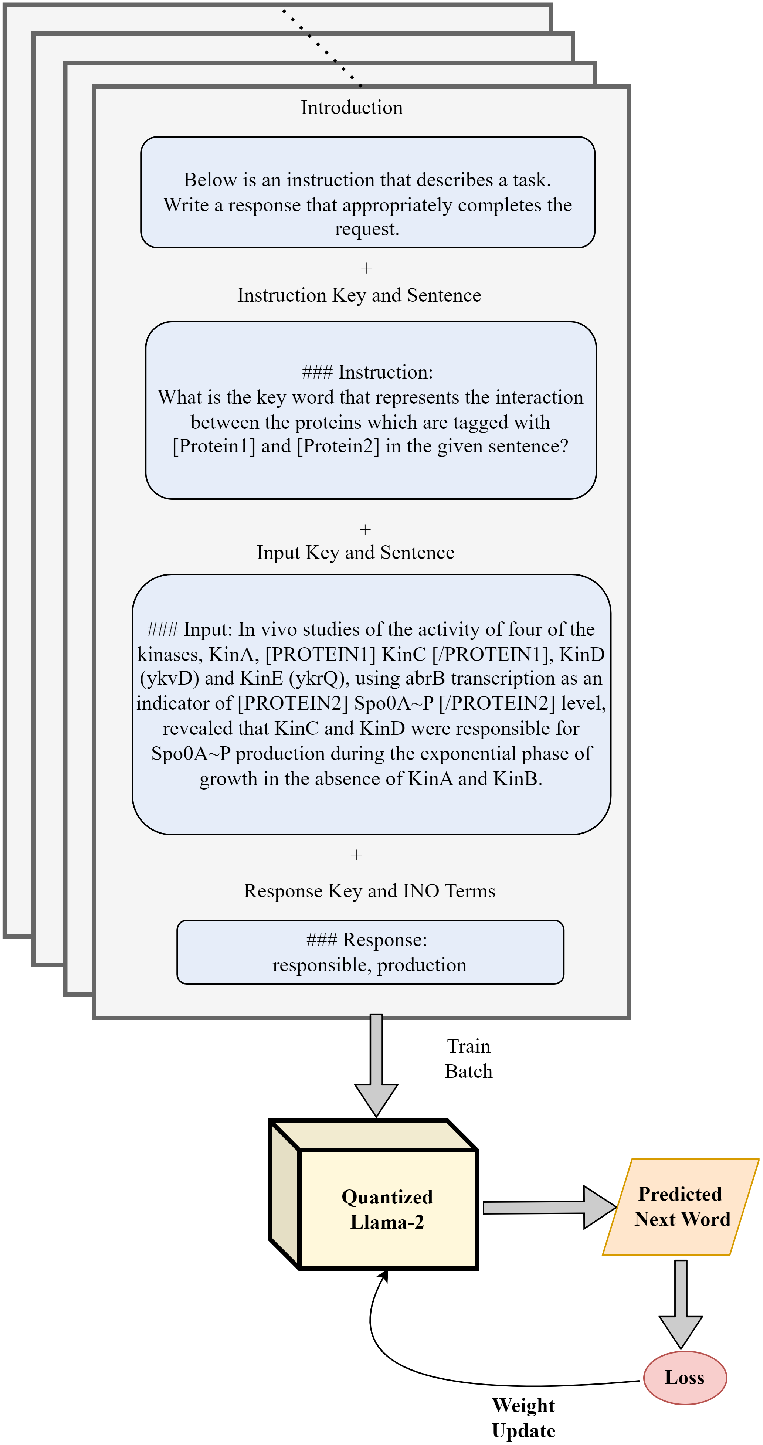
Instruction fine-tuning prompt example with a prompt that contains the introduction, instruction, input, and response statements. The quantized Llama-2 model is fine-tuned using next-word prediction.

## V. Experiments and Results

While evaluating the results of identifying relation keywords, we used the token-wise precision, recall, and F1 metrics. In computing these metrics, we first processed the predicted keywords by mapping them to the corresponding INO IDs ^2 3^ and performed the evaluation based on the predicted and actual INO IDs. Therefore, the INO IDs were used to standardize the evaluation of the result of the LLMs. Some examples of token-wise score computation are given in Table I.

**TABLE I.** Some examples of token-wise precision (p), recall (r), and f1 percentage calculations.

| Actual Keywords | Actual INO IDs | Predicted Keywords | Predicted INO IDs | P | R | F1 |
| --- | --- | --- | --- | --- | --- | --- |
| dependent, drives, expression, promoter | INO_0000157, INO_0000157, INO_0000102, INO_T000008 | dependent, promoter | INO_0000157, INO_T000008 | 1 | 0.5 | 0.67 |
| gene expression, activation | INO_0000102, INO_0000024 | gene expression, action | INO_0000102, INO_0000024 | 1 | 1 | 1 |
| converting | INO_0000058 | derivation | INO_0000058 | 1 | 1 | 1 |
| protein activation | INO_0000026 | protein activation by mutant | INO_0000028 | 0 | 0 | 0 |

### A. Results of In-Context Learning

#### 1) Zero-Shot Setting

In the zero-shot in-context learning setting, the test examples from the LLL INO dataset are given one by one to the Llama-2 models.

In Table II, the zero-shot in-context learning performance of Llama-2-7b-chat, Llama-2-13b-chat, and Llama-2-70b-chat on the test split of the LLL INO dataset are given. Since the Llama-2-70b-chat model is the largest parameterized model, it yields the highest F1-score, as expected. The Llama-2-13b-chat model achieved the second highest F1-score, trailing by 14.3%. Llama-2-7b-chat achieved the lowest F1-score, trailing by 41.2%.

**TABLE II.**
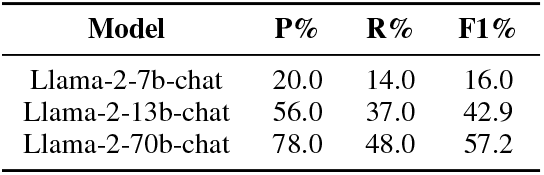
Zero-shot prompting inference scores as precision (P), recall (R), and F1.

| Model | P% | R% | F1% |
| --- | --- | --- | --- |
| Llama-2-7b-chat | 20.0 | 14.0 | 16.0 |
| Llama-2-13b-chat | 56.0 | 37.0 | 42.9 |
| Llama-2-70b-chat | 78.0 | 48.0 | 57.2 |

The Llama-2-7b-chat model generated responses that deviated far from the expected format. The Llama-2-70b chat model also responded with long paragraphs justifying the responses. This shows that the zero-shot in-context learning setting is not sufficient to use LLMs directly.

#### 2) Few-Shot Setting

In few-shot learning, the selection of context examples is not trivial. We performed three different experiments with different context examples selected from the LLL INO training split. The reported results were again on the test split.

In Table III, the performances of the Llama-2 chat models are given for three different experiments where the context examples are different. In the first experiment, the first forty samples from the training split of the LLL INO dataset are given as context examples. Similarly, the second and the third set of context examples are the subsequent forty samples in the second and third experiments.

**TABLE III.** Few-shot prompting inference scores as precision (P), recall (R), and F1 with different sets of context examples.

|  | Experiment 1 |  |  | Experiment 2 |  |  | Experiment 3 |  |  | Average |  |  |
| --- | --- | --- | --- | --- | --- | --- | --- | --- | --- | --- | --- | --- |
| Model | P(%) | R(%) | F1(%) | P(%) | R(%) | F1(%) | P(%) | R(%) | F1(%) | P(%) | R(%) | F1 (%) |
| Llama-2-7b-chat | 56.0 | 45.0 | 46.9 | 68.7 | 54.0 | 58.5 | 70.0 | 82.0 | 72.7 | 64.9 | 60.3 | 59.4 |
| Llama-2-13b-chat | 68.0 | 62.0 | 62.7 | 67.7 | 55.0 | 58.9 | 62.7 | 76.0 | 66.4 | 66.1 | 64.3 | 62.7 |
| Llama-2-70b-chat | 72.0 | 64.0 | 64.7 | 62.0 | 57.0 | 57.3 | 61.7 | 84.0 | 68.5 | 65.2 | 68.3 | 63.5 |

In “Experiment 1” given in Table III, the zero-shot setting of the Llama-2-70b-chat resulted in the highest F1-score. The Llama-2-13b-chat achieved the second highest F1-score, with a gap of 2%. The Llama-2-7b-chat achieved the lowest F1-score with a difference of 17.8%. The results of the fewshot setting with the second set of context examples are given in Table III as “Experiment 2”. The Llama-2-13b-chat achieved the highest F1-score, followed by the Llama-2-7b-chat. Interestingly, the Llama-2-70b-chat lags behind the 7 billion and 13 billion parameterized models by 1.2% and 1.6%, respectively. The Llama-2-7b-chat achieved the highest F1-score in Experiment 3 given in Table III compared to the Llama-2-13b-chat and Llama-2-70b-chat. The Llama-2-70b fall lag behind by 4.2%, and the Llama-2-13b fall lag behind by 6.3%. As revealed by the experiments, the gap between the scores of the individual models in each experiment is much smaller compared to the gap in the zero-shot setting. This indicates that the few-shot setting performs better on the unseen tasks. According to the averaged scores given as “Averaged” in Table III, the Llama-2-70b-chat achieved the highest F1-score overall. As expected, the Llama-2-13b-chat was in second place, while the Llama-2-7b-chat came last.

#### B. Results of Parameter-Efficient Instruction Fine-Tuning

We experimented with Llama-2-7B-chat and Llama-2-13B-chat models. Due to limited resources, we could not finetune the Llama-2-70b-chat. For the fine-tuning, we used the training split. The validation split was used to tune the hyperparameters. For the training epoch we tried 2, 4, 5, and 8. For the gradient accumulation steps we tried the default value, 1, and 4. For the learning rate, we investigated the 2e-4, 2e-5, 5e-4, and 5e-5. For the LoRa *α* and R parameters, we tried combinations of 4, 8, 16, 32, and 64. The tuned hyperparameters for training and the LoRa weights are given in Table IV. We also applied the early stopping strategy with patience as two on the validation loss.

**TABLE IV.** Hyper-parameters.

| Training Hyper-parameters | Value |
| --- | --- |
| Epoch for Llama-2-7b-chat | 5 |
| Epoch for Llama-2-13b-chat | 4 |
| Batch Size per Device | 1 |
| Gradient Accumulation Steps | 4 |
| Learning Rate | 2e-4 |
| LoRa Hyper-parameters | Value |
| $\alpha$ | 32 |
| R | 8 |

The training results of the individual Llama-2 chat models on test split are shown in Table V. The results demonstrate that the Llama-2-7b-chat and Llama-2-13b-chat performed similarly after fine-tuning. The Llama-2-13b-chat was more precise in its predictions, while the Llama-2-7b-chat predicted more of the true interaction keywords. Overall, the Llama-2-13b-chat outperformed the Llama-2-7b-chat by 1.1% on F1-score. Table VI presents some responses generated by the best-performing model, the fine-tuned Llama-2-13b-chat, from the LLL INO test split.

**TABLE V.** Llama-2 fine-tune scores as precision (P), recall (R), and F1.

| Model | P % | R % | F1 % |
| --- | --- | --- | --- |
| Llama-2-7b-chat | 79.3 | 84.0 | 80.2 |
| Llama-2-13b-chat | 86.0 | 80.0 | 81.3 |

**TABLE VI.** Some example responses of the fine-tuned Llama-2-13B-Chat.

| Sentence IDs | Keywords | Predicted Keywords | Sentence |
| --- | --- | --- | --- |
| 19 | dependent, expression | dependent | Therefore, the physiological role of [PROTEIN1] sigmaB [/PROTEIN1] -dependent [PROTEIN2] katX [/PROTEIN2] expression remains obscure. |
| 51 | dephosphorylation | dependent | Dephosphorylation of [PROTEIN2] SpoIIAA-P [/PROTEIN2] by [PROTEIN1] SpoIIE [/PROTEIN1] is strictly dependent on the presence of the bivalent metal ions Mn2+ or Mg2+ |
| 115 | recognize, promoter | recognize, promoter | We overproduced and purified sigma(X) from Escherichia coli and demonstrate that in vitro, both sigma(A) and [PROTEIN1] sigma(X) [/PROTEIN1] holoenzymes recognize promoter elements within the sigX-[PROTEIN2] ypuN [/PROTEIN2] control region. |
| 131 | dependent, drives, expression, | promoter dependent, promoter | The sigmaB-dependent promoter drives expression of yvyD under stress conditions and after glucose starvation, whereas a [PROTEIN1] sigmaH [/PROTEIN1]-dependent promoter is responsible for [PROTEIN2] yvyD [/PROTEIN2] transcription following amino acid limitation |
| 135 | dependent, inducible | dependent, induced | In addition to the typical [PROTEIN1] sigmaB [/PROTEIN1] -dependent, stress- and starvation- inducible pattern, [PROTEIN2] yvyD [/PROTEIN2] is also induced in response to amino acid depletion. |

## VI. Conclusion

We investigated the problem of interaction keyword identification by leveraging the power of the LLMs by applying in-context learning and parameter-efficient instruction finetuning. We have contributed to the literature by investigating the detection of INO relation keywords using LLMs. As far as we know, this study is the first of its kind. We believe that our work can help researchers further analyze biomedical relations by mapping the relation keywords into the ontological IDs.

In the real world, it is not feasible to utilize the largest model for every task due to its huge memory requirements and the inference latency proportional to the model size. Our goal was to show performance improvement by applying the in-context learning and fine-tuning methods. We showed that the finetuned Llama-2-7b-chat and Llama-2-13b-chat outperformed the Llama-2-70b-chat in in-context learning. The Llama-2-13b-chat achieved the highest F1-score in fine-tuning and outperformed the Llama-2-70b-chat in in-context learning with zero-shot and few-shot in F1-score by 24.1% and 17.8%, respectively.

We have demonstrated the effectiveness of using LLMs to explain protein-protein interactions by mapping the predicted interaction keywords to the INO ontology. The proposed approach can be used to explain relations between other types of biomedical entities using ontologies and LLMs.

Future work in this area will focus on more deeply integrating the INO ontology to improve the models’ performance further. Specifically, the hierarchical structure and semantic relationships defined in INO could be leveraged to construct more informative prompts for the language models. This may involve incorporating INO class definitions and interclass relationships directly into the prompts, providing the models with richer contextual information about the types of interactions and their connections. Additionally, the INO synonyms and term variants could be used to expand the set of keywords the models are trained to recognize, going beyond the specific terms in the training data. As the context length of LLMs increases, it will be possible to feed the models with a larger set of INO keywords using these prompting strategies, which we believe will be helpful in increasing the models’ performance. Furthermore, experimenting with different opensource LLMs could also yield valuable insights into future research directions.

## VII. Availability of data and materials

The LLL INO dataset is available at https://huggingface.co/datasets/bengisucam/LLLINO-tagged. The INO dictionary and the codes used in the current study are available in our GitHub repository https://github.com/bengisucam/Ontology-based-PPI-explanation-using-LLMs.

## Acknowledgment

The study was supported by the U.S. National Institute of Allergy and Infectious Disease (U24AI171008 to Y.H. and J.H.), and A.Ö. was partially supported by the GEBIP Award of the Turkish Academy of Sciences.

## Footnotes

1 The INO term has the IRI: http://purl.obolibrary.org/obo/INO0000117.

2 The INO ontology is available on: http://purl.obolibrary.org/obo/ino.owl

3 The INO GitHub repository: https://github.com/INO-ontology/ino

